# Calcium transient was triggered by Na^+^ influx in phase 0 and regulated by Ca^2+^ influx in phase 2 of the action potential in adult rat left ventricular cardiomyocyte

**DOI:** 10.1101/2025.10.15.682715

**Authors:** Long Chen, Yi-li Yang

## Abstract

**Background:** Ca^2+^-induced Ca^2+^ release (CICR) has been the prevailing model in cardiomyocytes for over 50 years. However, whether Na^+^ influx plays a direct role in triggering Ca^2+^ release has remained unclear. We previously discovered that action potential (AP) phase 0 triggered Ca^2+^ release while phase 2 regulated the decay phase during the Ca^2+^ transient (CT). We hypothesized that CICR might only exist under experimental conditions, not during physiological AP cycles.

**Methods:** Simultaneous recordings of ion currents and CT from the same rat left ventricular cardiomyocyte were employed to investigate the temporal relationship between cation influxes and CT.

**Results:** The reactivation time for CT was approximately 250 ms. AP phase 0 triggered the CT, while phase 2 only regulated CT width. Pulses from -40 to +80 mV with 6 ms duration were applied to validate whether the reverse Na^+^/Ca^2+^ exchanger (NCX) could trigger CT. Pulses of -40 and -30 mV with 6 ms duration induced giant Na^+^ influx and tail current, but only Na^+^ influx triggered CT, not tail current. Pulses from -20 mV to +80 mV with 6 ms duration triggered CT via tail current, not via reduced Na^+^ influx. In Ca^2+^ free solution, Na^+^ influx was still able to trigger CT until sarcoplasmic reticulum (SR) Ca^2+^ content was remarkably reduced. Moreover, increased reverse NCX current induced by ramp-like depolarizing pulses did not trigger CT but widened its shape. Additionally, when SR function was destroyed by caffeine (20 mM/L), field stimulations caused progressively elevated cytosolic Ca^2+^ levels, indicating that SR has a buffering function for maintaining cytosolic Ca^2^ concentration.

**Conclusions:** Once Na^+^ influx triggers CT, Ca^2+^ influx cannot trigger a second CT during an AP cycle due to the limitation imposed by the reactivation time. Na^+^ influx might enter SR via an unknown channel or transporter at the terminal cisternae during the AP and induce Ca^2+^ release by acting on the SR luminal-facing site of the ryanodine receptor 2 (RyR2) channel and/or by increasing the positive potential gradient across the SR membrane.

## 1 Introduction

During excitation-contraction coupling, whole-cell CT is crucial for transferring electrical activity mediated by the AP to mechanical contraction via contractile proteins. CT consists of rising phase and a decay phase. The rising phase is mediated by rapid Ca^2+^ release via RyR2 from the sarcoplasmic reticulum (SR, the specialized term for endoplasmic reticulum in myocytes). The decay phase is mainly mediated by Ca^2+^ reuptake via sarcoplasmic/endoplasmic reticulum Ca^2+^-ATPase 2a (SERCA2a) into the SR and by Ca^2+^ efflux via NCX and plasma membrane Ca^2+^-ATPase (PMCA) to the extracellular space. Compared to the decay phase, the mechanism of the rising phase is more controversial and has attracted greater attention.

CICR has been the subject of intense study in excitable cells for more than 50 years. CICR in cardiac myocytes has been reviewed in several standout articles focusing on CICR-related functions^(1)^, structures^(2)^, and both^(3)^. The discovery of Ca^2+^ release from SR via RyR2 in skinned frog cardiac muscle was first proposed over 50 years ago in 1972^(4,5)^, following the discovery of CICR in skinned frog skeletal muscle in 1970^(6,7)^. Later, CICR in intact cardiac myocytes was also confirmed^(8,9)^.

The fundamental hypothesis of CICR in cardiomyocytes posits that when Ca^2+^ levels between the SR and the sarcolemma (the specialized term for plasma membrane in myocytes) reach a threshold level, Ca^2+^ activates the cytosolic-facing site of RyR2 to induce Ca^2+^ release in a positive feedback manner. The consensus supporting CICR is mainly based on the following evidence: (1) CICR did not occur in cardiomyocytes in the absence of extracellular Ca^2+^ or when L-type Ca^2+^ current was blocked^(10,11)^. (2) Elimination of Na^+^ influx by Li^+^ or tetrodotoxin did not block CICR^(11)^. (3) Ca^2+^ influx, not depolarized voltage, triggered CT^(12)^. Depolarization across the sarcolemma to the Ca^2+^ reversal potential resulted in suppression of CT^(13)^. (4) The dyadic space (the region bounded by the t-tubule and SR) contains only ryanodine receptors, L-type Ca^2+^ channels, and calsequestrin, thus forming the cardiac dyad that is fundamental for CICR^(14)^.

Although consensus for CICR in cardiomyocytes has been reached, CICR appears self-contradictory with many unanswered questions. CICR is a positive feedback system in which Ca^2+^ release might become regenerative and activate even more distant ryanodine receptors until SR Ca^2+^ content decreases to a certain level, regardless of the magnitude of the trigger. In fact, Ca^2+^ release peak from RyR2 is graded and proportional to the peak amplitude of L-type current. This paradox led to the formation of the “local control theory”, indicating that each L-type Ca^2+^ channel and its functionally associated ryanodine receptor constitute a release unit, and whole-cell CT is an ensemble of all activated units. Many studies on this theory have not reached consensus, and further studies on the function and structure of the release unit are needed. Due to the positive feedback characteristic, the mechanism responsible for controlling SR release termination remains unclear, although several mechanisms that might contribute to RyR2 closure have been proposed without consolidated experimental confirmation^(15)^.

Beyond these controversial issues, studies on CICR have paid less attention to the relationship between AP kinetics and RyR2 function. Moreover, CICR theory is not linked to the effects of ion channels and/or transporters across the SR membrane, which are important in regulating SR Ca^2+^ release^(16)^. Additionally, it has been reported that CT could be triggered by membrane depolarization voltage, known as “the membrane voltage-sensitive calcium release mechanism” which coexists with CICR^(17,18)^. However, this voltage-triggered CT was attributed to contaminating Ca^2+^ influx^(19,20)^.

The effect of Na^+^ influx on triggering CT was intensively investigated in the early 1990s without reaching consensus. Na^+^ influx-induced CT has been explained by mixed Ca^2+^ influx via (1) Na^+^ channels, (2) Ca^2+^ channels, and (3) reverse NCX, due to loss of voltage control from large Na^+^ influx. It was reported that Li^+^ current carried by Na^+^ channels was unable to trigger CT, suggesting that the indirect effect of NCX, not Ca^2+^ influx via Na^+^ or Ca^2+^ channels, was involved in mediating CT^(21)^. However, experiments by another group demonstrated paradoxical results showing that Li^+^ current did not prevent CT under the same experimental conditions^(11)^. Moreover, very low colocalization coefficients between NCX and RyRs in rat ventricle indicated that NCX did not form part of the dyad in ventricular myocytes to trigger CT^(14)^.

In this study, we investigated the independent effect of Na^+^ influx on CT and ruled out the triggering effect of contaminating Ca^2+^ influx on CT during AP phase 0. By investigating the reactivation time for CT, we found that once Na^+^ influx in AP phase 0 triggered Ca^2+^ release from RyR2, CICR during AP phase 2 might not exist within an AP cycle. CICR might only be present under experimental conditions.

It should be noted that “influx” as described in this study refers to the inward current of cation.

## 2 Materials and Methods

### 2.1 Ethical approval

All animal experiments were approved by the Animal Care and Use Committee of the International Centre for Genetic Engineering and Biotechnology, Taizhou Region Research Centre (approval number 202410A014), whose policies adhere to the National Institutes of Health USA Guide for the Care and Use of Laboratory Animals (National Institutes of Health Publication No. 85-23, revised 2011). Every effort was made to minimize animal pain and distress.

### 2.2 Animals

Male adult Sprague-Dawley rats (300-350 g) from the Laboratory Animal Centre of the International Centre for Genetic Engineering and Biotechnology, Taizhou Region Research Centre, were used in this study. All rats received food and water ad libitum and were maintained at controlled temperature (22°C) with a constant 12 -hour light/ 12-hour dark cycle.

### 2.3 Chemicals

All chemicals for Tyrode’s and KB solutions and reagents were purchased from Sigma-Aldrich (USA).

### 2.4 Isolation of cardiac myocytes

The rat left ventricular myocytes were obtained by enzymatic dissociation. Following thoracotomy, hearts were quickly excised, mounted on a Langendorff apparatus and retrogradely perfused. The excised whole heart was first perfused at 37 °C with the HEPES-buffered solution containing (in mM/L) NaCl 137, KCl 5.4, NaH_2_PO_4_ 0.33, MgCl_2_ 1, HEPES 20, glucose 10, and gassed with 95% O_2_ and 5% CO_2_ (pH 7.4 with NaOH). The heart was then perfused with the same buffer with the additions of collagenase type II (Worthington Biochemical Corp, NJ, USA) (0.7 mg/ml) and CaCl_2_ (0.5 μM/L) for 40 minutes. Following removal of both atria and the right ventricle, the left ventricular myocytes were gently separated with forceps in the KB solution containing (in mM/L) L-glutamic acid 50, KCl 40, Taurine 20, MgCl_2_ 1, KOH 70, EGTA 0.5 HEPES 10, glucose 10 (pH 7.4 with KOH) without collagenase. The separated left ventricular myocytes were filtered and then kept in KB solution for 2 hours.

### 2.5 The CT recording from rat left ventricular myocytes by field stimulation and patch clamp

Adult rat left ventricular myocytes were loaded with the membrane-permeable acetoxymethyl ester form of the fluorescent Ca^2+^ indicator Fluo-4 AM (5 µM/L) (Dojindo Laboratories, Japan) for 30 minutes at 37°C. Cardiomyocytes with sharp coordinated fluorescence sparkling and contraction were selected by field stimulation with two platinum electrodes (20 V) by visual inspection. The region of interest was selected with an adjustable window. CT amplitudes were calculated as the difference between peak and diastolic fluorescent values after subtracting background fluorescence.

For field stimulation, electrical pacing (20 V, alternating polarities) at different frequencies was employed according to specific experimental requirements. For patch clamp-induced CT recording, intact ventricular myocytes were voltage- or current-clamped (Axopatch 200B, Molecular Devices Co., USA) using whole-cell ruptured-patch configuration. Patch clamp stimulation schemes are described in the corresponding Results sections.

The extracellular solution for field stimulation or patch clamp-induced CT contained (in mM/L): NaCl 137, KCl 5.4, NaH_2_PO_4_ 0.33, MgCl_2_ 1, CaCl_2_ 1.8, HEPES 20, glucose 10 (pH 7.4 with NaOH). For Na^+^-free extracellular solution, NaCl (137 mM/L) was replaced by choline-Cl (137 mM/L). The pipette solution for patch clamp stimulation contained (in mM/L): KCl 115, NaCl 20, EGTA 10, HEPES 10, glucose 5, K_2_ATP 3.0, TrisGTP 0.5 (pH7.2 with KOH). All experiments were carried out at room temperature (20-22°C). The number of replicates is indicated with each experiment. Each replicate represents a cell from a different animal.

## 3 Results

### 3.1 CT was triggered by AP phase 0

Current clamp configuration was used to trigger CT from rat left ventricular myocyte loaded with fluorescent dye (Fluo-4). The patch clamp amplifier controlled by Clampex (Molecular Devices, CA, USA) triggered sampling start of CT to ensure that AP and CT activities were recorded simultaneously. Results indicated that AP phase 0, which is mediated mainly by Na^+^ influx, triggered the CT. AP peak appeared earlier than the CT peak. AP phase 2, which is mediated mainly by Ca^2+^ influx, did not induce secondary CT as shown in Figure 1. This experimental result was confirmed in two additional rat left ventricular myocytes, in agreement with a previous study^(22)^.

**Figure 1:**
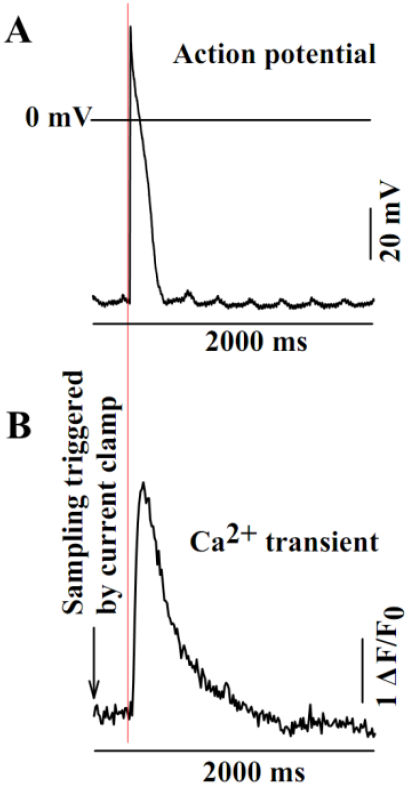
AP phase 0 mediated by Na^+^ dominated influx triggered the CT, not phase 2 mediated by Ca^2+^-dominated influx. (A) AP and (B) corresponding CT recorded simultaneously from the same enzymatically dissociated rat left ventricular myocyte. The arrow indicates the start of CT sampling, synchronized with AP recording. Horizontal bars indicate time scales; vertical bars indicate voltage magnitude (A) or fluorescence intensity (B).

### 3.2 Reactivation time for CT was regulated by isoproterenol (ISO)

To simulate the temporal sequence of Na^+^- and Ca^2+^-dominated influxes in the AP, a test pulse of 0 mV followed by a conditioning pulse of -40 mV was employed. The reactivation time for CT was approximately 250 ms (Figure 2D). Since the duration between the peak of phase 0 and the end of phase 2 in an AP is less than 100 ms, phase 2 is unable to trigger a second CT. Therefore, if CICR were valid, it could only occur in phase 0 mediated by Ca^2+^ influx mixed with giant Na+ influx, not in phase 2 dominated by Ca^2+^ influx.

**Figure 2:**
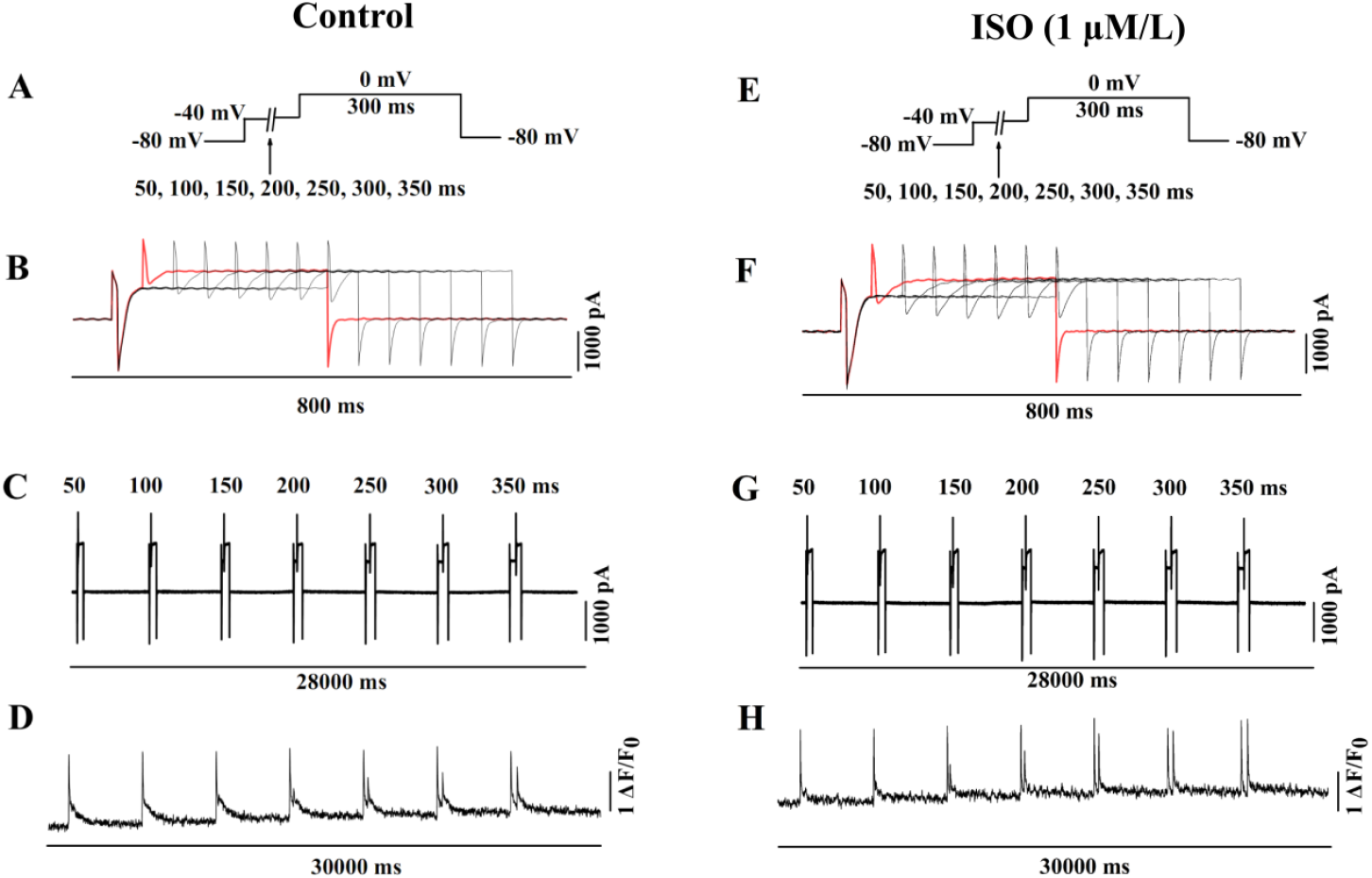
Isoproterenol (ISO, 1 μM/L) shortens the reactivation time for CT. Conditioning pulses of -40 mV with different durations preceded by test pulses of 0 mV for 300 ms were applied to the same myocyte. (A, E) Stimulation schemes for inducing Na^+^- and Ca^2+^-dominated influxes. (B, F) Selected portions from seven current traces. (C, G) Seven continuous traces with capacitances. (D, H) CTs corresponding to their stimulations. Red traces mark the first trace with 50 ms duration in panels B and F. Horizontal bars indicate different time scales; vertical bars indicate current magnitudes or fluorescence intensity.

Considering that isolated rat left ventricular myocytes used in these experiments lacked adrenergic agonists from neurohumoral regulation, isoproterenol (ISO) was employed to analyze the reactivation time for CT. The reactivation time was approximately 150 ms after ISO (1 µM/L) treatment (Figure 2H). ISO had minimal effect on Na^+^-dominated influxes induced by -40 mV pulses (Figure 2B, F) but significantly increased Ca^2+^-dominated influxes. The second Ca^2+^-dominated influx corresponding to the 100 ms pulse in Figure 2F, which could not trigger CT reactivation, was larger than any influx in Figure 2B. Therefore, CT reactivation might mainly depend on phosphorylated RyR2 by ISO, reflecting the intrinsic property of RyR2 release. This experimental result was confirmed in two additional rat left ventricular myocytes from different rats.

### 3.3 Effects of Na^+^-dominated influx and tail current on triggering the CT

To identify the minimal activation time for Na^+^-dominated influx while maximally reducing possible reverse NCX current and Ca^2+^-dominated influx, depolarizing pulses of -40 mV with different durations were employed. To identify the temporal relationship between current and CT, a time marker was generated by randomly activating the ZAP function (used to rupture the plasmalemma beneath the pipette tip; 0.5 ms stimulation pulse) on the Axopatch 200B amplifier to induce inward current and CT simultaneously.

As shown in Figure 3A, the 2 ms pulse did not induce Na^+^-dominated influx, tail current, or corresponding CT. The 4 ms pulse induced a confluent peak from Na^+^-dominated influx and tail current, along with corresponding CT. All pulses of 6, 8, 10, 12, and 14 ms induced Na^+^-dominated influxes and tail currents that were separated from each other. Peaks of Na^+^-dominated influxes were fixed at approximately the 3rd ms, and tail currents appeared at the end of pulses. Results indicated that pulses of -40 mV with ≥4 ms duration and time markers triggered corresponding CTs (Figure 3C), but tail currents did not induce CT due to limitation by reactivation time (Figure 3D).

**Figure 3:**
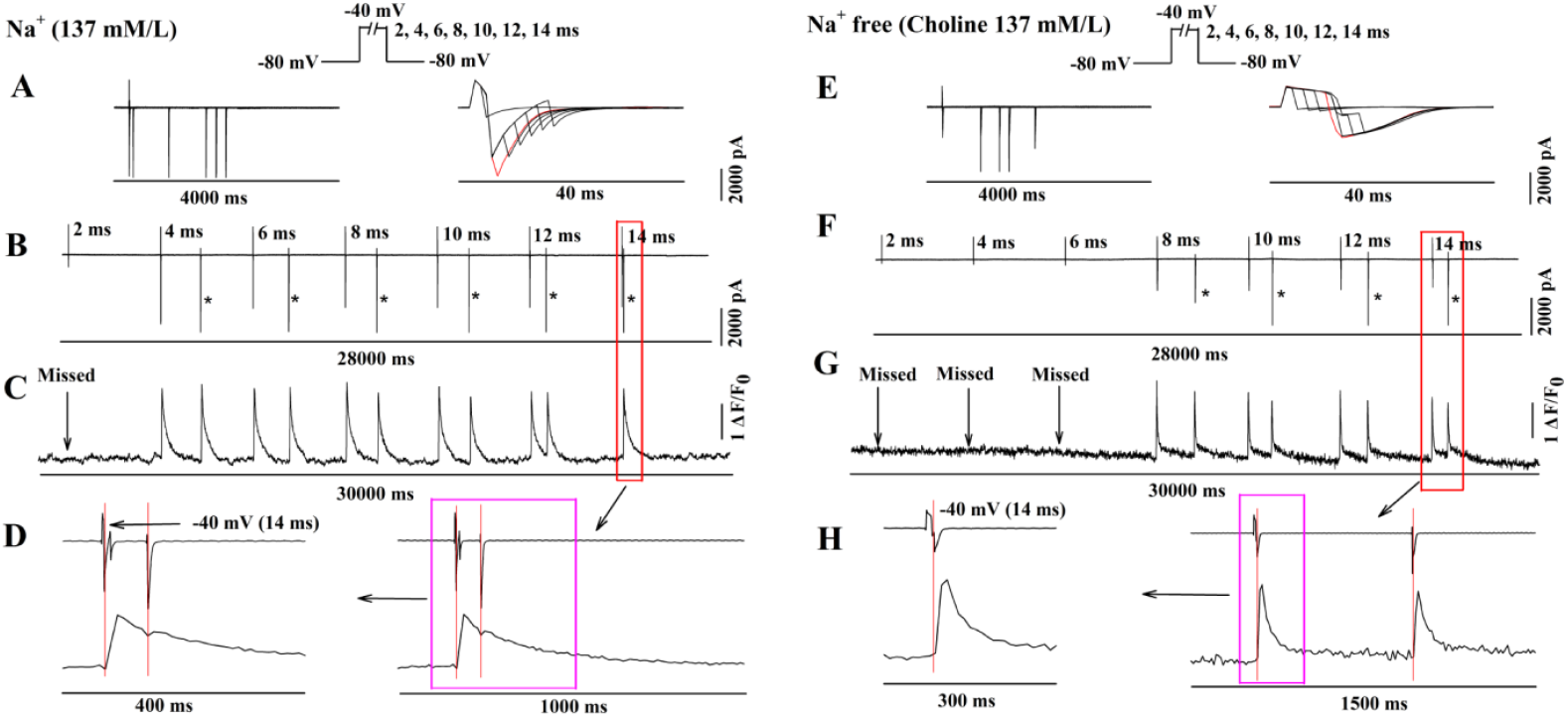
Na^+^-dominated influx, not tail current, triggers CT in normal Na^+^ solution. In Na^+^-free solution, tail currents trigger CT. Pulses of -40 mV with different durations were applied. Top panels show Na^+^-dominated current stimulation schemes. (A, E) Seven current traces and selected portions containing Na^+^-dominated currents and/or tail currents at different time scales. (B, F) Continuous seven current traces. (C, G) Corresponding CTs induced by Na^+^-dominated influxes, tail currents, and/or time markers. Red rectangles cover Na^+^-dominated influxes and tail currents induced by 14 ms pulses and time markers (B, F) with corresponding CTs (C, G). Red rectangles are expanded on the right in panels D and H. Scarlet rectangles in D and H are further expanded to analyze temporal correspondence between CT and Na^+^-dominated influx or tail current. “*” in panels B and F indicate capacitances and downward currents from time markers. Vertical arrows indicate missed CTs (C, G). Red traces mark confluent Na^+^-dominated influx and tail current peak induced by 4 ms pulse (A) and first tail current by 8 ms pulse (E). Horizontal bars indicate time scales; vertical bars indicate current magnitudes or fluorescence intensities.

To test the triggering effect of tail current on CT, Na^+^-free extracellular solution (replaced with choline) was used in another myocyte. The experimental protocol was identical to that in normal Na^+^ solution. No pulses induced Na^+^-dominated influx. Tail currents, which were slowly activated and inactivated, were only induced by pulses ≥8 ms duration. These tail currents induced by 8, 10, 12, or 14 ms duration triggered their corresponding CTs.

Combining these data, results induced by depolarizing pulses of -40 mV led to the following conclusions: (1) The peak of Na^+^-dominated influx appeared at the 3rd ms after the start of the -40 mV depolarizing pulse, and CT was triggered by Na^+^-dominated influx, not by tail current, in normal Na^+^ concentration solution. (2) No Na^+^-dominated influx appeared, and tail current was induced only by pulses ≥8 ms duration in Na^+^-free solution. (3) Tail currents exhibited fast activation and fast inactivation in normal Na^+^ solution, and slow activation and slow inactivation in Na^+^-free solution. (4) Ca^2+^ influx dominated the tail current in Na^+^-free solution. (5) Pulses of -40 mV with 2, 4, or 6 ms duration in Na^+^-free solution did not trigger CT, indicating no Ca^2+^ influx involvement via Na^+^ channels, Ca^2+^ channels, or reverse NCX. This experimental result was confirmed in two additional rat left ventricular myocytes from different rats.

### 3.4 Reverse NCX via voltage escape could not trigger CT

It was assumed that giant Na^+^ influx induced Ca^2+^ influx via reverse NCX through voltage escape. To ensure the independent effect of Na^+^ influx and rule out reverse NCX participation in triggering CT, pulses from -40 to +80 mV with 6 ms duration were employed in normal Na^+^ solution (Figure 4). These stimulations would fully activate Na^+^ channels while avoiding Ca^2+^ channel activation, and also activate possible reverse NCX by voltage escape up to +80 mV.

**Figure 4:**
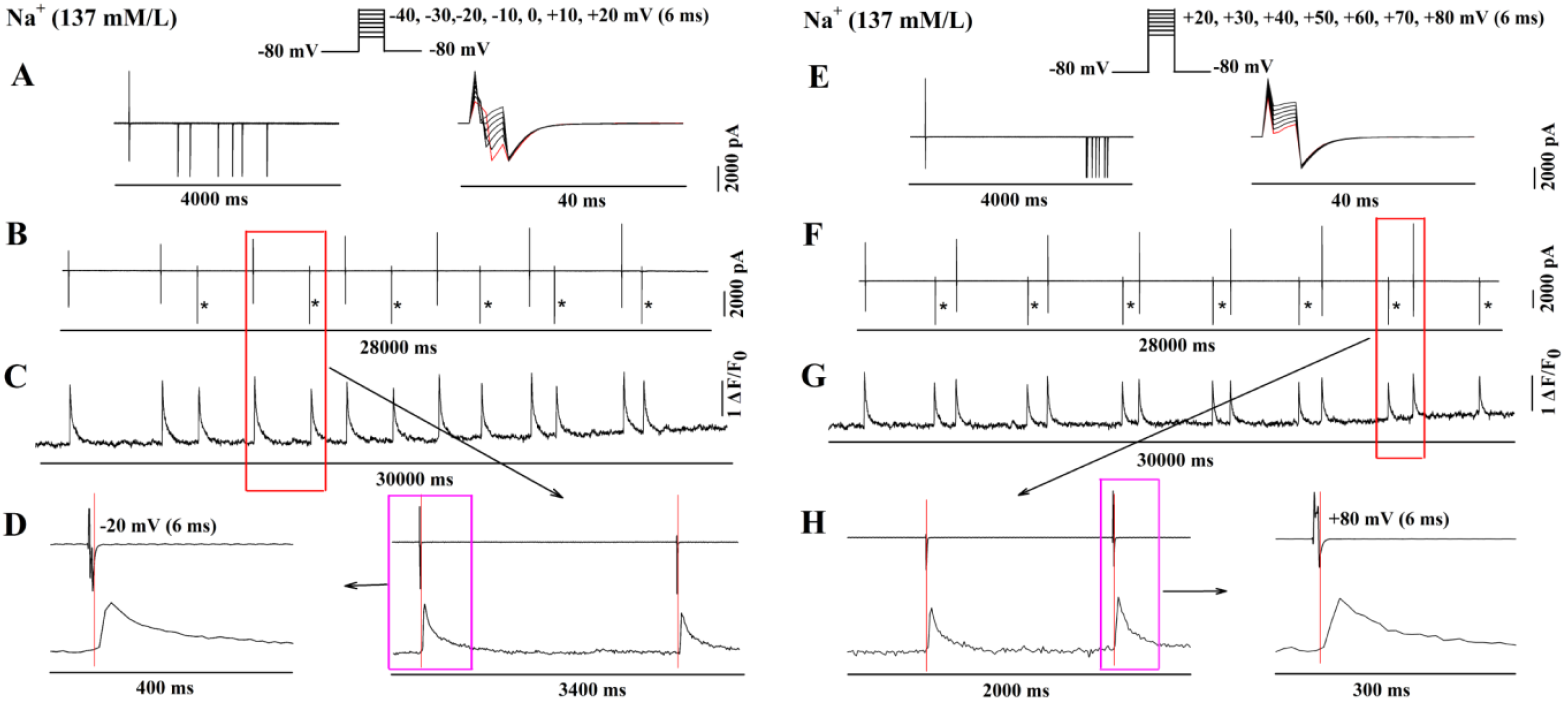
Giant Na^+^ influxes induced by -40 and -30 mV pulses trigger CT, while pulses from -20 to +80 mV trigger CT only via tail currents in normal Na^+^ solution. All pulses were 6 ms duration. Top panels show stimulation schemes. (A, E) Seven current traces and selected portions containing tail currents at different time scales. (B, F) Continuous seven current traces. (C, G) Corresponding CTs induced by Na^+^ influxes, tail currents, or time markers. Red rectangles cover Na^+^ influxes, tail currents, time markers (B, F) and corresponding CTs induced by -20 or +80 mV pulses (C, G). Red rectangles are expanded in D and H. Scarlet rectangles in D and H are further expanded to analyze temporal relationship between CT and Na^+^-dominated influx or tail current. “*” in B and F indicate capacitances and downward currents from time markers. Red traces mark currents induced by -40 or +20 mV pulses (A, E). Horizontal bars indicate time scales; vertical bars indicate current magnitudes or fluorescence intensities.

Pulses of -40 and -30 mV with 6 ms duration induced giant Na^+^ influx that triggered CT. However, induced Na^+^ influxes gradually reduced and disappeared with pulses from -20 to +80 mV. CTs were not triggered by pulses from -20 to +80 mV but were triggered by tail currents induced by their corresponding pulses (Figure 4D, H).

To further validate the independent effect of Na^+^ influx on triggering CT, Na^+^-free solution (replaced with choline) was applied (Figure 5). Pulses of -40 and -30 mV with 6 ms duration did not induce Na^+^ influx, tail current, or CT. Pulses from -20 mV to +80 mV induced consistent tail currents that were slowly activated and slowly inactivated. All CTs were triggered by tail currents only in Na^+^-free solution (Figure 5D, H).

**Figure 5:**
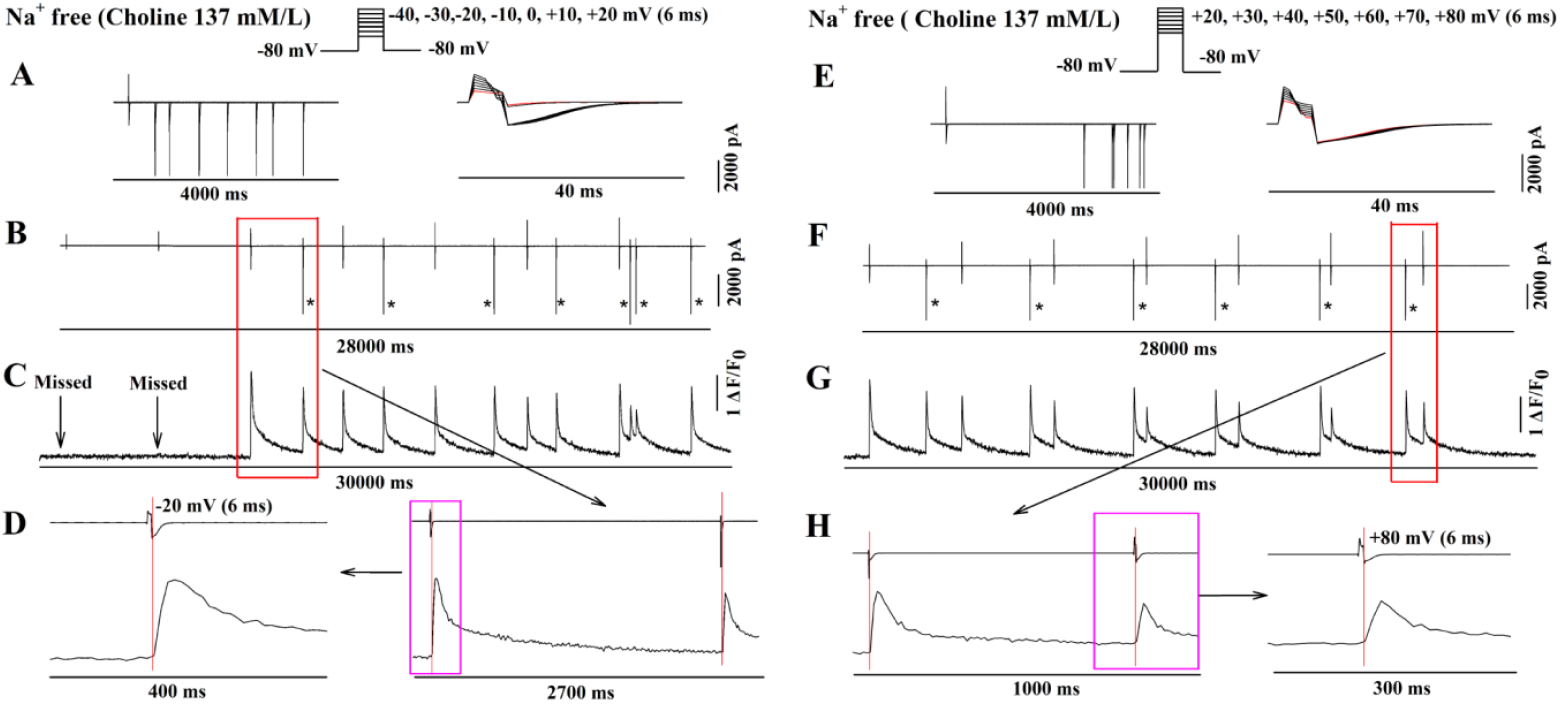
Pulses from -40 to +80 mV (6 ms duration) do not trigger CT in Na^+^-free solution. Tail currents induced by pulses from -20 to +80 mV and time markers trigger CTs. Top panels show stimulation schemes. (A, E) Seven current traces and selected portions containing tail currents at different time scales. (B, F) Continuous seven current traces. (C, G) Corresponding CTs induced by tail currents and time markers. Red rectangles cover tail currents, time markers (B, F) and corresponding CTs (C, G). Red rectangles are expanded in D and H. Scarlet rectangles in D and H are further expanded to analyze temporal relationship between CT and tail current. “*” in B and F indicate capacitances and downward currents from time markers. Red traces mark currents induced by -40 (A) or +20 mV (E) pulses. Horizontal bars indicate time scales; vertical bars indicate current magnitudes or fluorescence intensities.

Combining data from Figures 4 and 5, results indicated that CT was triggered by Na^+^ influx or tail current, but not by Ca^2+^ influx via Na^+^ channels, Ca^2+^ channels, or reverse NCX. Under normal physiological conditions, giant Na^+^ influx, not Ca^2+^ influx, triggered CT. Due to the limitation of reactivation time, a second CT could not be activated within an AP cycle. In other words, once CT was activated by a -40 mV pulse, Ca^2+^ influx via Ca^2+^ channels, reverse NCX, or tail current could not trigger a second CT. Voltage escape from giant Na^+^ influx might not exist, and Ca^2+^ influx via reverse NCX through voltage escape might be insignificant for triggering CT. This experimental result was confirmed in two additional rat left ventricular myocytes from different rats.

### 3.5 Relationships between Na^+^ and Ca^2+^ influxes, CT and SR Ca^2+^ content

Discontinuous field stimulation was used to induce CT for an extended period to explore how long CT could persist after extracellular Ca^2+^ was rapidly and completely eliminated. CT disappeared only after ≥5 minutes when normal extracellular Ca^2+^ (1.8 mM/L) solution was replaced with Ca^2+^-free solution. Moreover, SR Ca^2+^ content decreased dramatically as measured by caffeine (Figure 6A) compared to normal extracellular Ca^2+^ (1.8 mM/L) conditions (Figure 6C). However, CT persisted continuously in extracellular Ca^2+^ (1.8 mM/L) solution with unchanged configuration for >5 minutes (Figure 6D).

**Figure 6:**
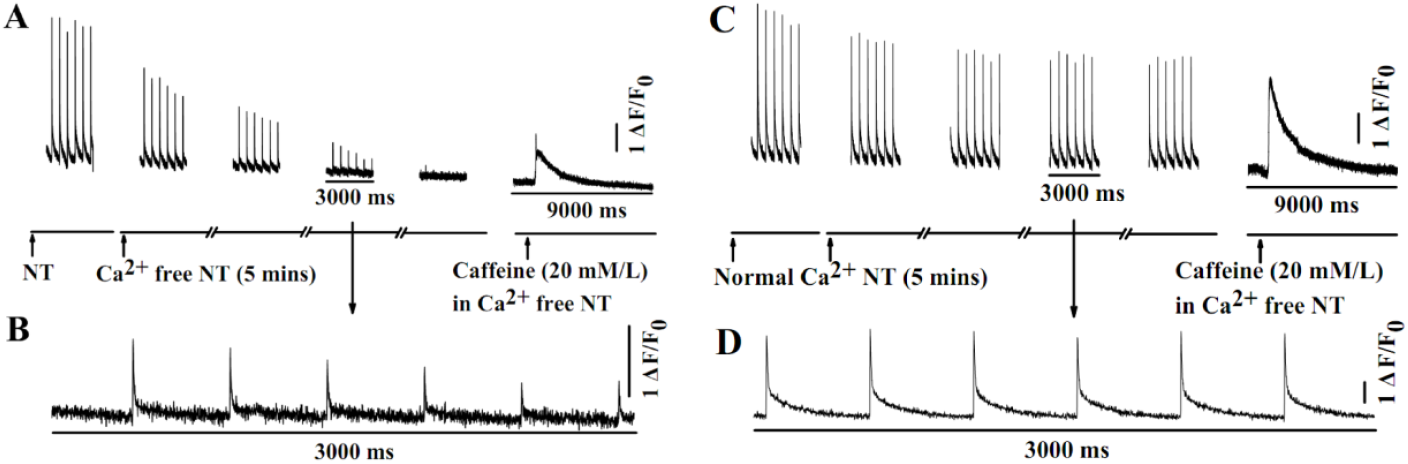
CT elimination and SR Ca^2+^ content reduction require > 5 minutes in Ca^2+^-free solution. Rat left ventricular cardiomyocytes were perfused with Ca^2+^-free normal Tyrode’s solution (NT) (A, B) or NT with Ca^2+^ (1.8 mM/L) (C, D) for 5 minutes with discontinuous field stimulation (20 V, 0.2 Hz). When field stimulation-induced CT disappeared after approximately 5 minutes of Ca^2+^-free NT perfusion, caffeine (20 mM/L) in Ca^2+^-free NT was rapidly applied (A). Similarly, caffeine (20 mM/L) in Ca^2+^-free NT was perfused to myocytes in Ca^2+^ (1.8 mM/L) solution after 5 minutes (C). Effects of Ca^2+^-free NT or normal NT on CT and SR content are displayed in A or C; corresponding expanded time scales of CTs are shown in B or D. Horizontal bars indicate time scales; vertical bars indicate fluorescence intensities.

These results indicated that the t-tubular system, which deeply penetrates the myocyte, constitutes a relatively sealed network forming a “Ca^2+^ micro-environment homeostasis in the t-tubule system”. Ca^2+^ in the t-tubule system would be replenished from subsarcolemmal space via NCX and PMCA until SR Ca^2+^ content was depleted in the presence of Ca^2+^-free extracellular solution. The CT shape in Ca^2+^-free solution resembled that of reverse Na^+^ current (Figure 6B) compared to that in Ca^2+^ solution (Figure 6D), indicating that CT kinetics by Na^+^ influx differed from that by Ca^2+^ influx. This experimental result was confirmed in three additional rat left ventricular myocytes.

To explore the effect of pure Na^+^ influx on triggering CT, a laborious experiment was performed in Ca^2+^-free solution. Currents and CTs were simultaneously recorded before and after extracellular Ca^2+^ was replaced with choline (137 mM/L) solution. Na^+^ and Ca^2+^ currents via patch clamp were applied to trigger corresponding CTs from the same myocyte simultaneously. During this experiment, Na^+^ current remained continuously unchanged while Ca^2+^ current gradually reduced until disappearing. Moreover, the ratio between CT amplitude induced by Na^+^-dominated influx and that by Ca^2+^-dominated influx increased gradually, as shown in Figure 7. Additionally, CT by Na^+^-dominated influx gradually transitioned to fast rising and fast decay configuration (from Figure 7D to G) compared to Figure 7A.

**Figure 7:**
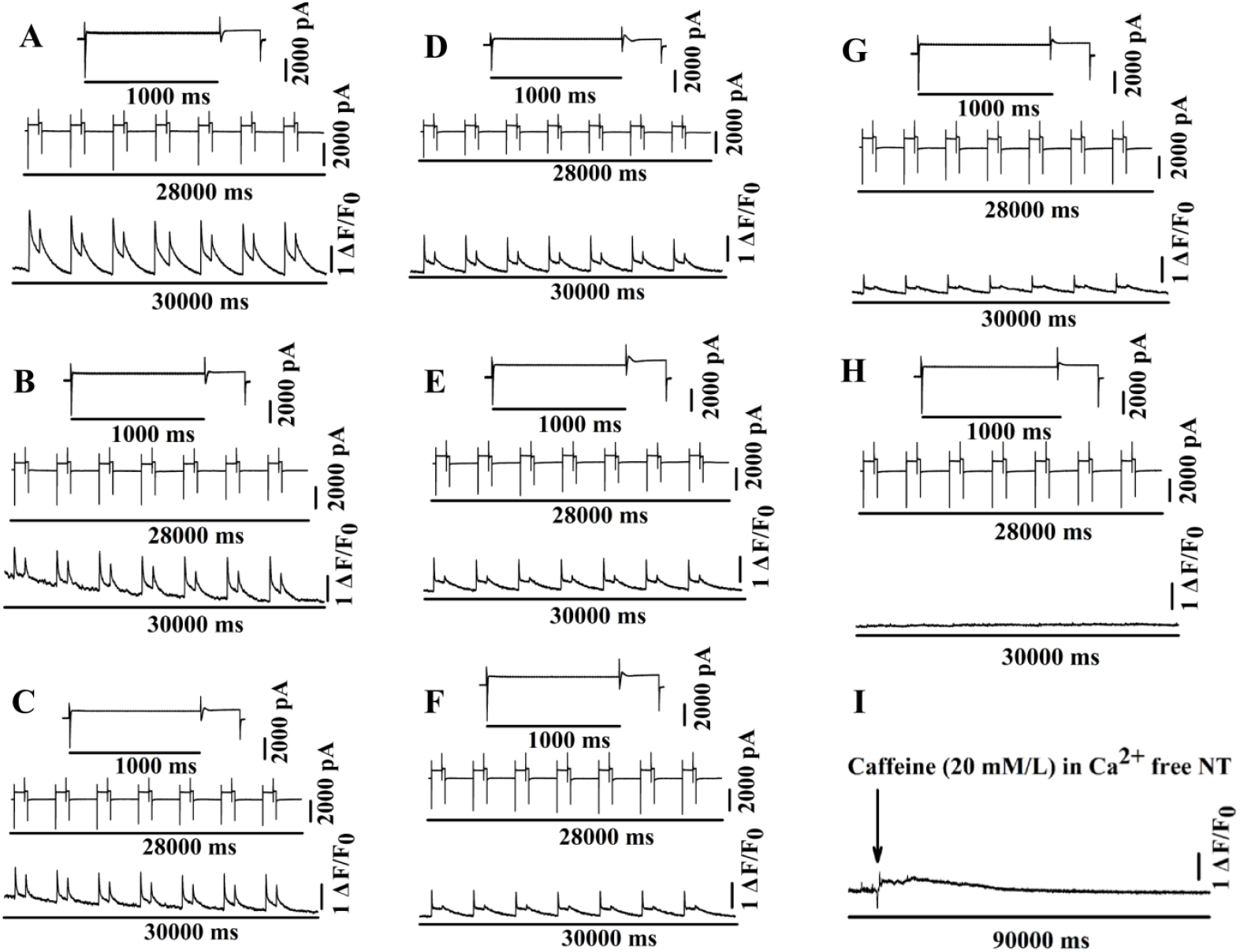
Effects of Ca^2+^-free extracellular solution on Na^+^- and Ca^2+^ -dominated currents, CT, and SR content. Upper panels show selected portions of seven Na^+^- and Ca^2+^-dominated current traces; middle panels show continuous seven current traces; lower panels show CTs. Panels A-H correspond to different time points (0, 30, 60, 90, 120, 150, 180, and 210 s) from the beginning of Ca^2+^ -free solution application. Once CT disappeared, SR content was measured by rapid application of caffeine (20 mM/L) in Ca^2+^ -free solution (I). Horizontal bars indicate time scales; vertical bars indicate current magnitudes or fluorescence intensity.

Once CT disappeared, caffeine (20 mM/L) in Ca^2+^-free solution was perfused to the recorded myocyte. SR content was dramatically reduced by extracellular Ca^2+^-free application in approximately 5 minutes. This result indicated that the failure of Na^+^ influx to trigger Ca^2+^ transient in Ca^2+^-free solution might be due to reduced SR Ca^2+^ content, not the ability of Na^+^ influx to trigger release. This experimental result was confirmed in two additional rat left ventricular myocytes.

To explore the effect of Na^+^-free and Ca^2+^-free conditions on triggering CT, choline (137 mM/L) was used to replace Na^+^. After application of Na^+^-free and Ca^2+^-free solution, Na^+^-dominated influx disappeared gradually while Ca^2+^-dominated influx persisted even when CT disappeared. Both CTs induced by reduced Na^+^- or Ca^2+^-dominated influxes decreased gradually. The ratio between CT amplitude induced by Na^+^-dominated influx and that by Ca^2+^-dominated influx decreased gradually, as shown in Figure 8. Once CT disappeared, caffeine (20 mM/L) in Na^+^- and Ca^2+^-free solution was used to investigate SR content. SR Ca^2+^ content diminished remarkably, and it took longer to reach the peak of caffeine-induced Ca^2+^ release from SR, indicating that lack of Na^+^ in extracellular solution resulted in slower Ca^2+^ expulsion from the cytosol. This experimental result was confirmed in two additional rat left ventricular myocytes.

**Figure 8:**
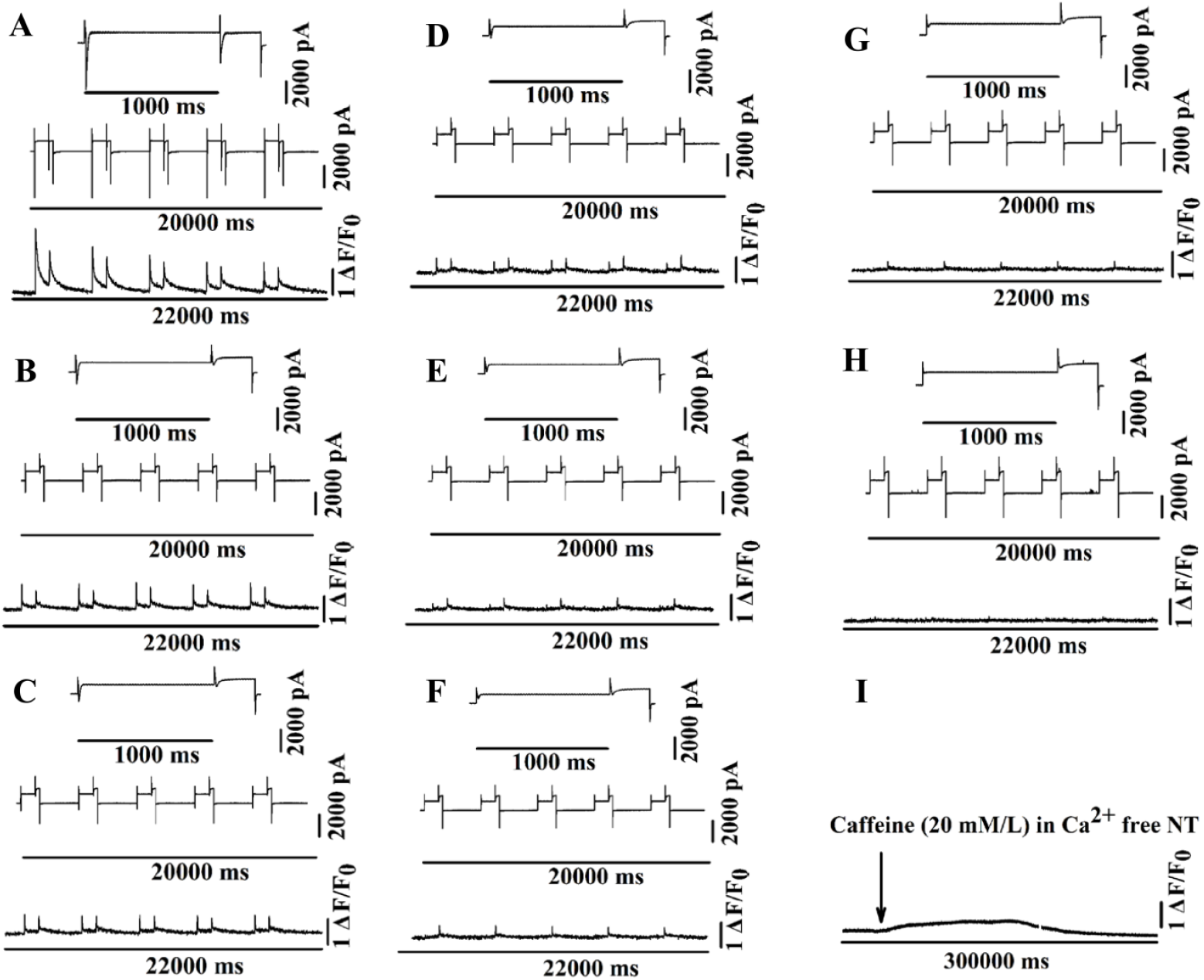
Effects of Na^+^-free and Ca^2+^-free solution on Na^+^- and Ca^2+^-dominated influxes, CT, and SR content. Upper panels show selected portions of seven Na^+^- and Ca^2+^-dominated influx traces; middle panels show continuous seven current traces; lower panels show CTs. Panels A-H correspond to different time points (0, 30, 60, 90, 120, 150, 180, and 210 s) from the beginning of Na^+^-free and Ca^2+^-free solution application. Once CT disappeared, SR content was measured by rapid application of caffeine (20mM/L) in Na^+^- and Ca^2+^-free solution (I). Horizontal bars indicate time scales; vertical bars indicate current magnitudes or fluorescence intensity.

### 3.6 Effect of reverse NCX current on widening CT shape

As strong evidence supporting CICR, CT induced by Na^+^-dominated influx has been triggered by Ca^2+^ influx via reverse NCX current during the AP. In this experiment, pharmacological and electrophysiological methods were used to explore the impact of reverse NCX current on CT. 4-Aminopyridine (4-AP, 2 mM/L), an I_to_ channel blocker, was used to inhibit I_to_ current, resulting in prolonged AP plateau and enhanced Ca^2+^ influx via reverse mode NCX. Results indicated that 4-AP did not enhance CT amplitude but widened CT shape compared to control (Figure 9A, B), in agreement with a previous study^(23)^.

**Figure 9:**
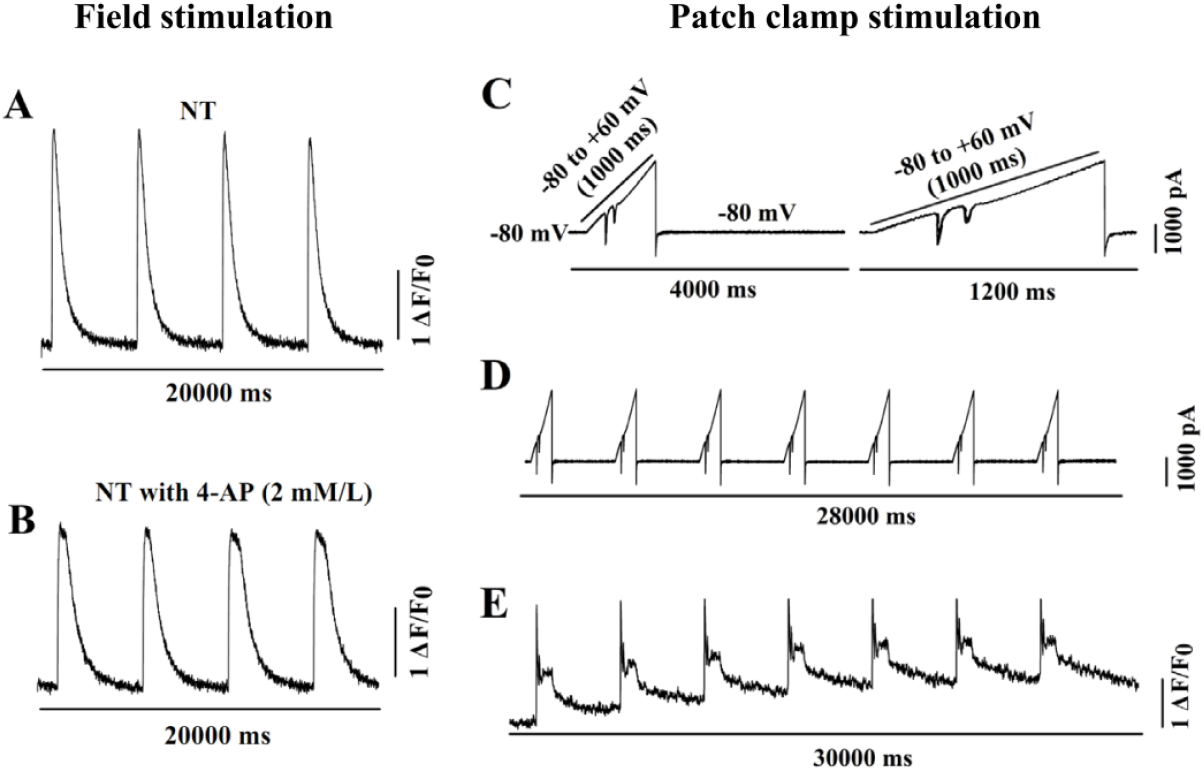
Reduced I_to_ current by 4-aminopyridine (4-AP, 2 mM/L) increases reverse NCX current, resulting in widened CT shape. Field stimulation (20 V, 0.5 Hz)-induced CTs before and after 4-AP (2 mM/L) are shown in A and B. (C) Seven current traces and selected portions of currents induced by ramp-like depolarizing test pulse. (D) Continuous seven current traces. (E) CTs corresponding to ramp-like pulses. Horizontal bars indicate time scales; vertical bars indicate current magnitudes or fluorescence intensities.

To further explore the impact of reverse NCX current, a ramp-like depolarizing pulse from a holding potential of -80 mV to +60 mV for 1000 ms was employed. This ramp-like pulse induced Na^+^- and Ca^2+^-dominated influxes at positions of 27% or 39% from the ramp-like line beginning, accounting for their activation voltages of -42 or -25 mV, respectively. Obviously, reverse NCX current gradually increased following the ramp-like pulse. The gradually increased reverse NCX current did not activate a second CT. However, it apparently widened CT shape and elevated diastolic cytosolic Ca^2+^ concentration. This experimental result was confirmed in two additional rat left ventricular myocytes.

### 3.7 Effect of SR on buffering cytosolic Ca^2+^ level

In this experiment, caffeine (20 mM/L) in normal Tyrode’s solution (NT) with Ca^2+^ (1.8 mM/L) was perfused. Caffeine caused giant Ca^2+^ release from SR with slow recovery back to normal cytosolic Ca^2+^ level, which was mediated by PMCA and NCX. In this case, CT would not be triggered by either Ca^2+^- or Na^+^-dominated influx. Once cytosolic Ca^2+^ concentration level reached its lowest point (Figure 10B), field stimulation was again applied to the myocyte to observe cytosolic Ca^2+^ level changes. Instead of CT, field stimulation induced a Ca^2+^ wave with shape similar to that of reverse Ca^2+^ current. Basal levels of Ca^2+^ waves were elevated gradually. In this case, NCX and PMCA had insufficient capacity to expel Ca^2+^ out of the myocyte to balance cytosolic Ca^2+^ level. Therefore, SR might have a buffering effect on cytosolic Ca^2+^ level, indicating that Ca^2+^ might enter SR to trigger CT via RyR2 by acting on the luminal-facing site of RyR2 across the SR during an AP cycle. This experimental result was confirmed in two additional rat left ventricular myocytes.

**Figure 10:**
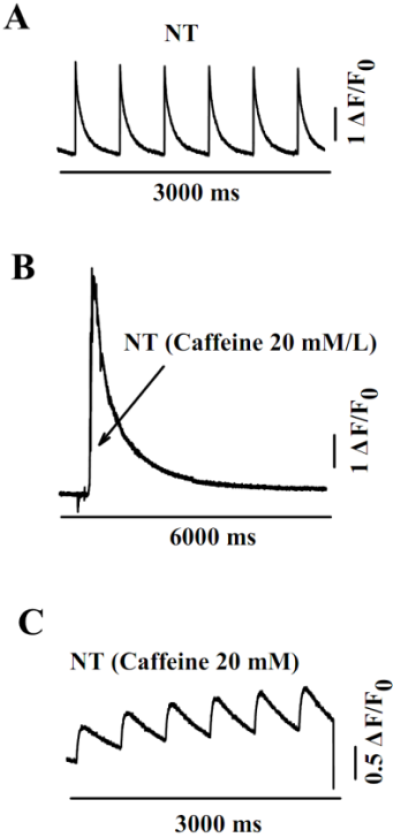
Cytosolic Ca^2+^ level changes induced by field stimulation after SR function is destroyed by caffeine (20 mM/L). (A) Field stimulation-induced CTs. (B) Caffeine (20 mM/L)-induced rapid giant Ca^2+^ release from SR with slow recovery. Arrow indicates the rising phase of Ca^2+^ release from SR by caffeine. (C) Ca^2+^ waves induced by field stimulation after caffeine treatment. Horizontal bars indicate time scales; vertical bars indicate fluorescence intensities.

## 4. Discussion

### Recordings of Na^+^-dominated and Ca^2+^ -dominated currents

For patch clamp recordings in this study, ion channel inhibitors were not used to isolate pure Na^+^ and Ca^2+^ currents to avoid any possible impacts on SR channels by these inhibitors. Due to the limited compensation range of the amplifier, the test potential of -40 mV to induce Na^+^ current could not be maintained consistently due to possible voltage escape. The recorded Na^+^ current was contaminated with possible Ca^2+^, I_to_, and reverse NCX currents. Ca^2+^ influx might enter the myocyte through Na^+^ channels via “slip-mode conductance”^(24)^. Therefore, Na^+^ current was termed Na^+^-dominated current in this study. Similarly, for recording Ca^2+^ current, 4-AP to block Ito and Cs^+^ to block K^+^ current were not used. The recorded Ca^2+^ current, which was contaminated with I_to_ and other types of K^+^ currents, was termed Ca^2+^-dominated current. In fact, the patch clamp technique is unable to isolate any pure current based on the non-selective properties of channels and lack of highly specific inhibitors. In this study, to investigate the triggering effect of pure Na^+^ influx, Ca^2+^-free extracellular solution was used. In this case, pure Na^+^ current was obtained by eliminating Ca^2+^ influx via Na^+^ channels, Ca^2+^ channels, and reverse NCX.

### Novel findings in this study

Voltage change during an AP in cardiomyocytes is transient, ranging from rapid depolarization up to +40 mV by Na^+^ influx to resting potential of -80 mV in a cascade manner through alternative activation and inactivation of all types of Na^+^, K^+^, and Ca^2+^ channels. Experimental technique limitations have resulted in the controversial issue of CICR remaining unresolved for many decades. For example, dynamic cytosolic and SR Ca^2+^ concentrations can hardly be measured simultaneously, and microstructures such as channels, transporters, and receptors across the SR membrane are poorly understood. Although the patch clamp technique was unable to mimic the transient voltage changes during an AP, the specific test potential corresponding to its current is powerful for investigating relationships between voltage, current, and CT in the same cardiomyocyte. As a result, several innovative findings in this study are summarized as follows:

#### (1) AP phase 0 triggered CT

AP recordings demonstrated that phase 0 triggered CT, not phase 2. However, massive Ca^2+^ influx in phase 2 might regulate CT width. Therefore, if CICR theory were valid, CICR might only occur in phase 0 mediated by Na^+^-dominated influx mixed with Ca^2+^ influx. The effect of peak Ca^2+^ influx via L-type Ca^2+^ channels on CICR might only exist under experimental conditions.

#### (2) Kinetics of CT

The reactivation time for CT was 250 ms and was shortened by β-adrenergic agonist to 150 ms, indicating that Ca^2+^ reactivation depends not only on the magnitude of Na^+^ or Ca^2+^ influx but also on RyR2 intrinsic properties. The reactivation time ensures that AP triggers CT only once, and electrical arrhythmia might not induce contractile arrhythmia. In fact, the anesthetized rat heart rate may reach 450 beats per minute, accounting for a CT cycle of 143 ms without any contractile disorder. The effects of ISO on shortening reactivation time and accelerating CT inactivation induced by Na^+^-dominated influx might explain this controversial issue. The isolated rat left ventricular myocytes used in these experiments lacked adrenergic agonists from neurohumoral regulation. ISO modified CT kinetics most likely by regulating the activation and inactivation of Na^+^ and Ca^2+^ influxes and by modifying RyR2 intrinsic properties through phosphorylation.

#### (3) Reverse NCX current could not trigger CT

Except for Ca^2+^ current, the reverse mode of NCX is considered to play an important role in regulating CT during AP phase 0(^25, 26)^. The reversal potential of NCX depends on intracellular Na^+^ concentration, with more negative reversal potential in the presence of higher intracellular Na^+^ concentration^(26)^. In other words, massive Na^+^ entering the cytosol via Na^+^ channels during AP phase 0 results in more negative reversal potential and greater Ca^2+^ entry via reverse NCX. However, it has been reported that NCX is located far from RyR2 and has no impact on RyR2 release^(14, 27)^. NCX density in the dyad was less than that of L-type channels in atrial myocytes^(28)^.

Reverse NCX current might not directly trigger CT in AP phase 0 based on our data as follows: First, the -40 mV pulse with 6 ms duration induced giant Na^+^ influx and triggered CT. However, the +80 mV pulse with 6 ms duration to simulate voltage escape and maximally induce reverse NCX did not trigger CT; instead, CT was triggered by its tail current. Second, 4-AP, which inhibits I_to_ resulting in increased reverse NCX current, did not enhance CT amplitude but prolonged the CT peak. Third, the ramp-like test pulse when potential reached +60 mV did not induce CT reactivation. Reverse NCX might only regulate the width of the CT decay phase. Considering that the CT peak could be widened by Ca^2+^ influx or reverse NCX, Ca^2+^ release most likely has two types of release modes: one might be massive release called CT, and another might be small amounts of release called Ca^2+^ physiological leak.

#### (4) Ca^2+^ recycling between t-tubule and subsarcolemmal space in Ca^2+^-free extracellular solution

It is commonly assumed that when extracellular Ca^2+^ is quickly eliminated, Ca^2+^ current via L-type Ca^2+^ channels and CT would disappear and SR content would be exhausted immediately. Our data in this study indicated that Ca^2+^ current and CT gradually reduced and lasted up to 5 minutes. The t-tubular system, which deeply penetrates the myocyte, constitutes a relatively sealed network forming a “Ca^2+^ micro-environment homeostasis in the t-tubule system”. Ca^2+^ in the t-tubule system would be replenished from subsarcolemmal space via NCX and PMCA until SR Ca^2+^ content was depleted in the presence of Ca^2+^-free extracellular solution. Compared to t-tubular Ca^2+^ being relatively independent of extracellular Ca^2+^, Na^+^ was relatively dependent on extracellular Na^+^. Na^+^ current disappeared earlier than Ca^2+^ current in the presence of Na^+^-free and Ca^2+^-free solution, indicating that the reserve capacity of Ca^2+^ supported by SR in the cytosol was more powerful than that of Na^+^.

#### (5) Na^+^ influx independently triggered CT

Many efforts have been made to investigate whether Na^+^ influx could independently trigger CT. First, pure Na^+^ current using pharmacological and electrophysiological methods was used to support this hypothesis. However, pure Na^+^ current was difficult to isolate from Ca^2+^ influx via Ca^2+^ channels and reverse NCX, resulting in controversial conclusions about the triggering effect of Na^+^ influx. Second, Na^+^ influx in Ca^2+^-free solution failed to trigger CT, indicating that only Ca^2+^ influx was able to trigger CT, not Na^+^ influx. Third, Ca^2+^ influx in Na^+^-free solution could trigger CT, which further strengthened CICR theory.

In this study, we designed a series of experiments to verify the triggering effect of Na^+^ influx alone on CT. First, the -40 mV pulse with 4 ms duration, which could not activate Ca^2+^ channels, was able to trigger CT. Moreover, pulses from -20 to +80 mV with 6 ms duration, which gradually reduced Na^+^ influx, could not trigger Ca^2+^ transient, but their corresponding tail currents did, indicating that Ca^2+^ influx via Na^+^ channels, Ca^2+^ channels, and reverse NCX was unable to trigger CT with 6 ms pulse duration.

Second, to further verify that Na^+^ current was able to independently trigger CT without Ca^2+^ influx involvement, we continuously recorded Na^+^ and Ca^2+^ currents and CT from the same cardiomyocyte when Ca^2+^-free extracellular solution was rapidly and completely replaced. Results demonstrated that both CTs induced by Na^+^- or Ca^2+^-dominated influxes reduced gradually until SR content decreased remarkably. During application of Ca^2+^-free solution, Na^+^ current peak remained unchanged and Ca^2+^ current decreased gradually until disappearing. However, CT induced by Ca^2+^-dominated influx decreased proportionally more than that by Na^+^ current. In this way, Na^+^ current could not be mixed with Ca^2+^ influx via L-type Ca^2+^ channels or reverse NCX. This experiment demonstrated that Na^+^ influx was able to independently trigger CT.

Third, the fact that Na^+^ influx failed to trigger CT in Ca^2+^-free solution, as reported previously, might be due to dramatically decreased SR content, not due to Na^+^ influx itself.

Logically, triggering of CT should not be mediated by currents such as L-type Ca^2+^ current that are highly regulated by neurohumoral factors. Large and rapidly activated Na^+^ current, which is minimally regulated, is more suitable to trigger CT. The slowly activated Ca^2+^ current via L-type channels might be designed mainly to shape CT width according to sympathetic tone to adapt to cardiac output requirements. However, Ca^2+^ current is also important for continuously maintaining SR Ca^2+^ content and CT amplitude during AP phase 0.

### Possible CT mechanism during an AP cycle

Because of the early establishment of CICR theory, the consensus that Ca^2+^ through L-type Ca^2+^ channels targets the cytosolic-facing site of RyR2 has never been questioned. Two key issues about CICR termination and graded magnitudes of Ca^2+^ release relative to Ca^2+^ influx are poorly understood. Several mechanisms proposed to explain CICR termination remain unclear, such as time-dependent inactivation, stochastic attrition, allosteric coupling between RyRs, SR Ca^2+^-dependent RyR2 luminal gating changes, and pernicious attrition^(15)^.

The local control theory is used to explain the graded Ca^2+^ release issue based on the premise that Ca^2+^ influx spreads unevenly over the cardiomyocyte. In fact, whole-cell L-type Ca^2+^ current magnitude depends on the ensemble of all single-channel open probability and voltage-driven current amplitude rather than the number of activated channels scattered on the plasmalemma. In other words, all L-type Ca^2+^ channels are activated evenly during the AP. Any minimal Ca^2+^ influx that could trigger recorded CT would be the ensemble of several hundred activated single Ca^2+^ channel currents. It is difficult to imagine that a large number of activated Ca^2+^ channels spread only over a localized region. In addition, Ca^2+^ release from the terminal cisternae of the SR, which is far from troponin at an angle of 90 degrees, would not be efficient in energy utilization. Therefore, graded Ca^2+^ release by Ca^2+^ influx cannot be reasonably explained by “local control theory” which is key for CICR.

Taken together, Na^+^ influx might enter the SR lumen mediated by an unknown channel via terminal cisternae of the SR during AP phase 0 and act at the luminal-facing site sensor of RyR2 to open it. The rapid rise of positive potential in SR by massive Na^+^ influx forces rapid Ca^2+^ release from RyR2, termed CT. Meanwhile, an unknown transporter expels Na^+^ out of SR, and negative ions are taken into SR to terminate Ca^2+^ release. Expelled Na^+^ from SR is then transported outside the cell via NCX. During AP phase 2, Ca^2+^ might also enter the SR lumen to balance SR Ca^2+^ concentration and is released to the cytoplasm in leak mode due to RyR2 inactivation induced by previous Na^+^-triggered CT. We speculate that SR Ca^2+^ is released to a specific space where troponin is coupled to RyR2. It has been reported that SERCA2 distribution in the heart is surprisingly sparse and localized to the z-line^(29)^ and myofilaments^(30)^, indicating that reuptake by SERCA2a is localized near RyR2 release sites for minimum energy consumption. In this way, the CT decay phase might be mediated by (1) rapid termination of RyR2 Ca^2+^ release, (2) reuptake by SERCA2a, and (3) Ca^2+^ diffusion away from the troponin region. Moreover, Ca^2+^ channels located in t-tubules are embedded deeply into the cytoplasm and form a relatively sealed system that is relatively independent of extracellular Ca^2+^ concentration changes. We propose that Na^+^ or Ca^2+^ influx works as a trigger, and RyR2 as the release mechanism aimed at troponin for maximized utilization of SERCA2a.

### Conclusions

CT triggered by Na^+^ influx, which is minimally regulated, ensures consistent CT amplitude. Ca^2+^ current, which is regulated by neurohumoral tone, might control CT width to adapt to cardiac output requirements. The reactivation time for CT, which is longer than 150 ms, ensures that the heart rarely exhibits contractile arrhythmia even when electrical arrhythmia occurs. Moreover, the amount of Ca^2+^ reuptake by SERCA2a might be less than previously thought, based on the finding that Ca^2+^ influx directly enters SR to act at the luminal-facing site of RyR2 during an AP cycle.

## Abbreviations and Acronyms

AP: action potential
CICR: Ca^2+^ induced Ca^2+^ release
CT: Ca^2+^ transient
NCX: Na^+^/Ca^2+^ exchanger
RyR2: ryanodine receptor 2
PMCA: plasma membrane Ca^2+^-ATPase
SERCA2a: sarcoplasmic/endoplasmic reticulum Ca^2+^-ATPase 2a
SR: sarcoplasmic reticulum

## Author Contributions

L. Chen designed the research study and conducted experiments and data analysis. Y. L. Yang prepared and edited the manuscript.

